# Transcutaneous cervical vagus nerve stimulation improves sensory performance in humans

**DOI:** 10.1101/2023.08.08.552508

**Authors:** Michael Jigo, Jason B. Carmel, Qi Wang, Charles Rodenkirch

**Affiliations:** Sharper Sense, Inc., New York, NY; Department of Neurology and Orthopedics, Columbia University Medical Center, New York, NY; Department of Biomedical Engineering, Columbia University, New York, NY; The Jacobs Technion-Cornell Institute at Cornell Tech, New York, NY

## Abstract

Accurate senses depend on high-fidelity encoding by sensory receptors and error-free processing in the brain. Progress has been made towards restoring damaged sensory receptors. However, methods for on-demand treatment of impaired central sensory processing are scarce. Prior invasive studies demonstrated that continuous vagus nerve stimulation (VNS) in rodents can activate the locus coeruleus-norepinephrine system to rapidly improve central sensory processing. Here, we investigated whether transcutaneous VNS improves sensory performance in humans. We conducted three sham-controlled experiments, each with 12 neurotypical adults, that measured the effects of transcutaneous VNS on metrics of auditory and visual performance, and heart rate variability (HRV). Continuous stimulation was delivered to cervical (tcVNS) or auricular (taVNS) branches of the vagus nerve while participants performed psychophysics tasks or passively viewed a display. Relative to sham stimulation, tcVNS improved auditory performance by 37% (p=0.00052) and visual performance by 23% (p=0.038). Participants with lower performance during sham conditions experienced larger tcVNS-evoked improvements (p=0.0040). Lastly, tcVNS increased HRV during passive viewing, corroborating vagal engagement. No evidence for an effect of taVNS was observed. These findings validate the effectiveness of tcVNS in humans and position it as a method for on-demand interventions of impairments associated with central sensory processing dysfunction.

## Introduction

Accurate and detailed sensory perception is vital for navigating daily life. But cognitive constraints – including aging, inattention, and fatigue – reduce the precision with which the brain processes sensory stimuli and lead to consequential misperceptions^1–3^. Recent research in rodents showed that activation of the locus coeruleus-norepinephrine system (LC-NE), a major subcortical noradrenergic nucleus, can increase sensory processing fidelity when stimulated directly or indirectly via the vagus nerve^4–6^. In humans, branches of the vagus nerve traverse the neck and ear and project to the brain where it modulates the LC-NE via the nucleus tractus solitarius pathway^6–9^. Transcutaneous stimulation of these vagal branches is tolerated and safe^10^. In this study, we investigated the potential of transcutaneous vagus nerve stimulation (tVNS) for improving sensory performance in humans.

Neuromodulatory systems, such as the LC-NE, are well-known to modulate the functioning of subcortical and cortical sensory processing pathways^11–17^. Noradrenergic and cholinergic systems play influential roles in inducing attentive and alert behavioral states that promote heightened sensory performance^14–20^. Recent work in animal models has shown that during continuous activation of LC-NE, somatosensory processing is rapidly enhanced via NE-mediated suppression of thalamic burst spiking that would otherwise degrade detailed sensory transmission^4^. Continuous invasive vagus nerve stimulation in rodents has been shown to similarly activate the LC-NE to generate steady sensory enhancements that parallel direct LC activation. These NE-mediated sensory enhancements were found to have a rapid onset and be transient, occurring during ongoing stimulation and dissipating when stimulation ends^5^. Collectively, these findings demonstrate that direct activation of the LC-NE system, or indirect activation via vagus nerve stimulation, generates rapid improvements to central sensory processing that persist during stimulation delivery.

By coupling these recent neuroscientific advances with contemporary neuromodulation techniques, it may be possible to enhance human sensory performance safely and on demand. tVNS emerges as a promising approach due to its ability to safely activate the vagus nerve in humans^10^. Neuroimaging studies support this potential by demonstrating that tVNS can modulate brain activity in a manner consistent with vagus nerve activation^21–24^. Transcutaneous auricular VNS (taVNS) targets the auricular branch of the vagus nerve in the ear^21,23,25^, presumably a primarily afferent fiber due to its peripheral nature^25,26^, whereas transcutaneous cervical VNS (tcVNS) most likely targets both afferent and efferent fibers of the cervical branch in the neck^8,22,24^. Furthermore, stimulation of terminal vagal fibers that project into the laryngeal and pharyngeal muscles^27^ neighboring the tcVNS site, may also provide antidromic vagal activation during tcVNS. However, the reliability of tVNS in activating the LC-NE in humans, as assessed by non-invasive NE markers such as pupil diameter, P300, and salivary alpha amylase, has to this point been inconclusive^28^.

Here, we investigated the hypothesis that continuous tVNS improves metrics of sensory performance in neurotypical human adults. We further examined cardiac markers commonly used to demonstrate efferent vagal activation^28,29^. Using a crossover design, we conducted three experiments that included control conditions (no stimulation or continuous sham stimulation at the forearm, ear, or neck), and active taVNS or tcVNS conditions. We explored the efficacy of two different patterns of tcVNS (30 Hz and triple pulse) and taVNS (30 Hz and 3 Hz) that have been shown to drive LC activity in rodents during invasive VNS^30^ and are based on parameters commonly used for tVNS^28^. While receiving stimulation, participants passively viewed a computer display or performed auditory gap discrimination and visual letter discrimination psychophysics tasks that are constrained by central sensory processing.

Briefly, our findings reveal that tcVNS improved auditory performance by 37% and visual performance by 23%, on average, compared to control conditions. Notably, tcVNS led to greater improvements in individuals with poorer sensory performance during control conditions. Furthermore, tcVNS increased heart rate variability (HRV) during passive viewing, corroborating vagal engagement. However, tcVNS-evoked changes in HRV were absent during active task engagement. These results provide evidence that continuous tcVNS can engage the vagus nerve to improve sensory performance in humans and suggest more extensive testing is warranted to probe the clinical relevance of using tcVNS to induce NE-mediated enhancement of central sensory processing.

## Results

We investigated the impact of tcVNS, taVNS, and sham stimulation on sensory performance across three experiments (Figure 1A), two stimulation frequencies (30 Hz and 3 Hz), and two waveforms (single pulse – 30 Hz and 3 Hz stimulation – and triple pulse; Figure 1B). In total, 29 individuals participated in the study and each experiment had a sample of 12 participants. Continuous stimulation was delivered while participants performed sensory tasks constrained by central sensory processing, namely auditory gap discrimination and visual letter discrimination tasks (Figure 1C). Adaptive psychophysical procedures determined sensory thresholds in each task, representing the shortest gap duration or smallest letter size that yielded 75% perceptual accuracy (Figure 1D). Stimulation conditions were completed in a counterbalanced or randomized order, except in Experiment 2 where conditions were pseudo-randomized (Figure 1E). In Experiment 2, Baseline and Arm controls served as a conditioning period that familiarized participants with the stimulation sensation and sensory tasks, after which Sham and tVNS conditions were performed. To avoid potential confounds arising from this pseudo-randomization procedure, Baseline and Arm conditions from Experiment 2 were excluded from analyses. Sensory tasks (and passive viewing in Experiment 2) were performed during each testing block, with continuous stimulation delivered throughout.

**Figure 1.**
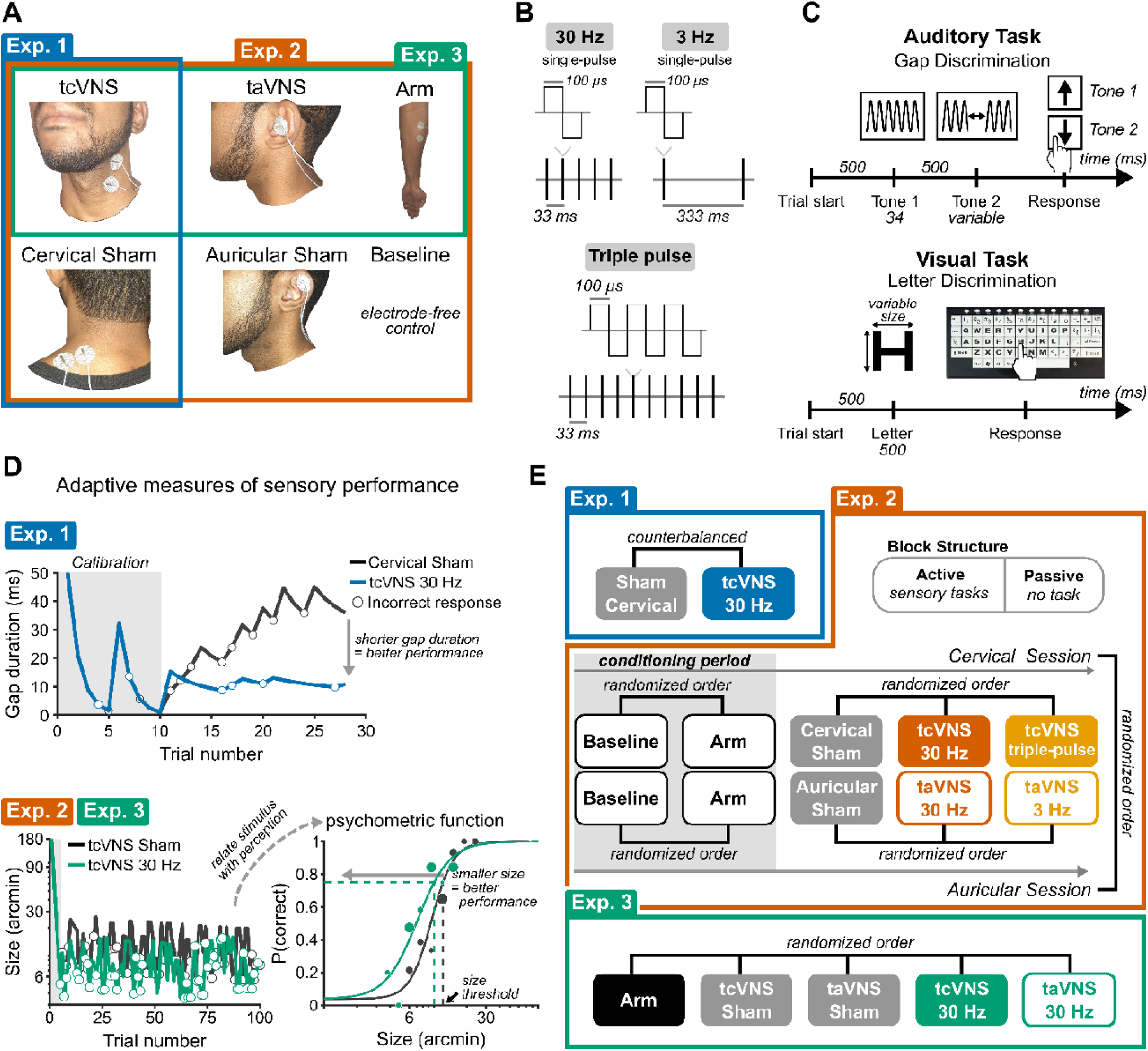
Stimulation protocols and psychophysics procedures. **(A)** Stimulation sites. Electrode placement shown on an author. Colored boxes group the stimulation sites tested in each experiment. **(B)** Stimulation waveforms and frequencies. **(C)** Trial sequence for the auditory and visual sensory tasks. (*top*) Trials of the auditory task consisted of two tone stimuli – one tone contained a gap and participants reported whether it occurred in Tone 1 or 2. (*bottom*) Trials of the visual task comprised a brief letter display and participants reported its identity using a keyboard. **(D)** Adaptive psychophysical procedures for measuring auditory (*top*) and visual sensory performance (*bottom*). The panels depict data from representative participants. In Experiment 1, the adaptive procedure initially presented gap durations that were easily judged correctly but became progressively more difficult until converging on a duration that maintained 75% accuracy. In Experiments 2 and 3, the adaptive procedure dynamically presented stimuli that yielded performance ranging from near-chance to near-perfect. Logistic functions were fit to trial-wise data and related perceptual accuracy with changes in the sensory stimulus; the size of individual dots on the logistic function depict the number of trials. An adaptive procedure for the visual task is displayed and a similar procedure was followed for the auditory task. The gray highlighted regions depict the initial sweep that calibrated the adaptive procedures. **(E)** Order of stimulation conditions in each experiment. Brackets indicate the randomization procedure. The inset in Experiment 2 labeled “Block structure” depicts the order in which participants completed sensory tasks and passive viewing within each stimulation block.

### tcVNS improved auditory performance

The gap discrimination task probed participants’ ability to detect silent gaps in sound, a critical function for decoding speech that depends on central auditory temporal processing^31,32^. Gap duration thresholds represented sensory performance during each stimulation condition, with shorter gap duration thresholds corresponding to heightened auditory performance. We delivered two tcVNS waveforms (single pulse and triple pulse; Figure 1B) continuously at 30 Hz while participants performed the task. Linear mixed models (LMM) determined the effectiveness of both tcVNS waveforms in all 29 participants across three experiments (n=120 observations; Figure 2A), finding a 37% improvement to auditory performance relative to control stimulation (p=0.00052, for a full LMM summary see Table S1).

**Figure 2.**
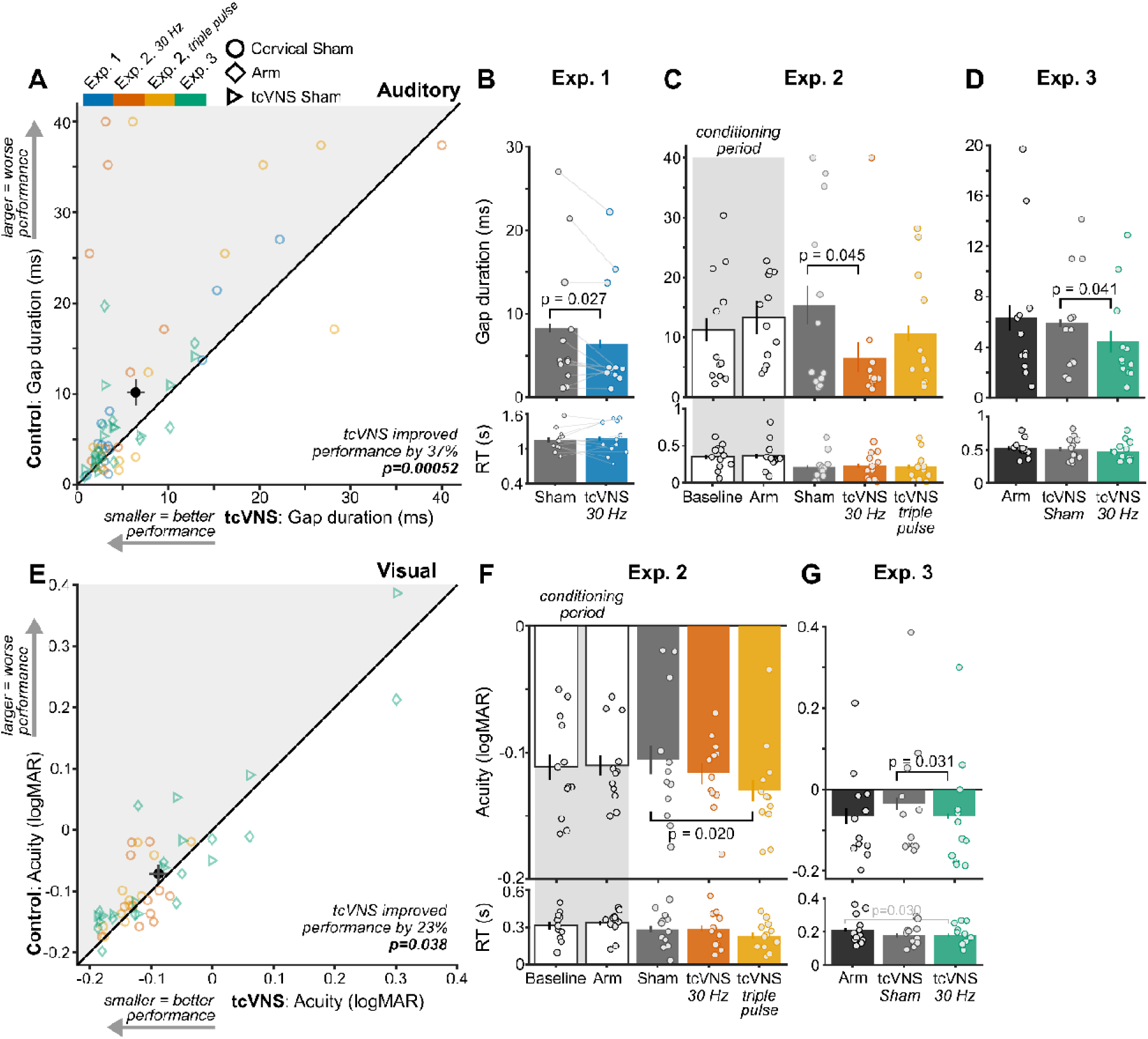
tcVNS improved sensory performance. **(A)** Gap duration thresholds during control conditions (Arm, Cervical Sham, tcVNS Sham) and active tcVNS conditions across all experiments and stimulation protocols (30 Hz and triple pulse). Each colored point depicts a participant in an experiment or stimulation protocol. The black point and error bars show the study-wide average and ±1 standard error of the mean (SEM) for tcVNS (horizontal line) and control conditions (vertical line). Points within the gray shaded region above the diagonal line indicate tcVNS-evoked sensory improvements. The p-value denotes the result of a linear mixed model that used the full dataset. **(B)** Gap duration thresholds in each stimulation condition of Experiment 1. Each point depicts an individual participant, bars depict the group-average, and error bars denote ±1 within-subject SEM^34^. The p-values denote results of a paired t-test; p-values for non-significant comparisons are not displayed. Subsequent bar plots follow these conventions. **(C)** Gap duration thresholds in each stimulation condition of Experiment 2 and **(D)** Experiment 3. The p-values denote results of permutation paired t-tests. **(E)** Visual acuity, in logMAR units, during control conditions and tcVNS conditions across experiments and stimulation protocols. Plotting conventions follow those in A. **(F)** Visual acuity thresholds in each stimulation condition of Experiment 2 and **(G)** Experiment 3.

tcVNS consistently evoked improvements within individual experiments that each comprised 12 participants (Figure 2B-D). tcVNS 30 Hz significantly improved performance relative to Cervical Sham in Experiment 1 (t(11)=2.54, p=0.027, d [95% CI] = 0.73 [0.072, 1.00]) and Experiment 2 (t(11)=2.32, p=0.045, d [95% CI] = 0.67 [0.29, 1.16]), and relative to tcVNS Sham in Experiment 3 (t(11)=2.36, p=0.041, d [95% CI] = 0.68 [0.17, 1.52]). However, performance during Arm stimulation in Experiment 3 did not differ from tcVNS 30 Hz (t(11)=1.58, p=0.16, d [95% CI] = 0.46 [-0.062, 1.05]). Unexpectedly, tcVNS triple pulse did not significantly improve performance relative to Cervical Sham in Experiment 2 (t(11)=0.68, p=0.54, d [95% CI] = 0.20 [-0.42, 0.84]). This difference among tcVNS protocols potentially arose from the small sample sizes in individual experiments. However, study-wide analyses that benefited from heightened statistical power found that tcVNS improved performance despite differences among stimulation waveforms (Figure 2A). Together, these findings demonstrated robust tcVNS-evoked improvements that occurred in both study-wide analyses and reliably across multiple controls (tcVNS Sham and Cervical Sham).

Improvements to auditory performance were not mediated by speed-accuracy tradeoffs – the propensity to make quick but error-prone or slow but accurate responses^33^. Participants’ reaction times (RTs) did not differ significantly among conditions in which tcVNS improved performance, namely between tcVNS 30 Hz and Cervical Sham in Experiment 1 (p=0.67) and Experiment 2 (p=0.86), and tcVNS Sham in Experiment 3 (p=0.64).

### tcVNS improved visual performance

The visual task probed participants’ ability to distinguish among small letters, a common measure of visual acuity that depends on central visual processing^35,36^. Letter size thresholds were transformed to logMAR units, a clinical measure of visual acuity^36^, with smaller logMAR values corresponding to heightened visual performance. We delivered two tcVNS waveforms (single pulse and triple pulse; Figure 1B) continuously at 30 Hz while participants performed this visual task. LMMs assessed the combined effectiveness of both tcVNS waveforms in all participants across Experiments 2 and 3 (n=96 observations; Figure 2E), finding a 23% improvement to visual performance relative to control stimulation (p=0.038, Table S2).

tcVNS consistently evoked improvements within both experiments, each comprising 12 participants (Figure 2F-G). In Experiment 2, tcVNS triple pulse significantly improved performance relative to Cervical Sham (t(11)=2.81, p=0.020, d [95% CI] = 0.81 [0.56, 1.42]). In Experiment 3, tcVNS 30 Hz significantly improved performance relative to tcVNS Sham (t(11)=2.53, p=0.031, d [95% CI] = 0.73 [0.20, 1.71]), but not Arm stimulation (t(11)=0.021, p=0.99, d [95% CI] = 0.0060 [-0.51, 0.81]). Unexpectedly, tcVNS 30 Hz did not significantly improve performance relative to Cervical Sham in Experiment 2 (t(11)=0.67, p=0.56, d [95% CI] = 0.19 [-0.58, 0.74]). Like the auditory results, these differences among tcVNS protocols potentially arose from the small sample sizes in individual experiments. Nonetheless, study-wide analyses found robust tcVNS-evoked effects despite differences among stimulation waveforms (Figure 2E).

Visual improvements were not mediated by speed-accuracy tradeoffs. RTs did not significantly differ among conditions that exhibited tcVNS-evoked improvements, namely between tcVNS triple pulse and Cervical Sham in Experiment 2 (p=0.33) and between tcVNS single-pulse and tcVNS Sham in Experiment 3 (p=0.83).

### tcVNS-evoked improvements scaled with sensory performance

We leveraged the natural variation in participants’ sensory performance during control conditions to examine whether it predicted the magnitude of tVNS-evoked sensory improvements (Figure 3). Regression revealed positive slopes between sensory performance during controls and the percent change in performance evoked by tcVNS (n=108 observations; Figure 3A, Table S3) and taVNS (n=96 observations; Figure 3B, Table S4). We examined whether regression to the mean, a statistical phenomenon that can generate false positives in repeated measures experimental designs^37^, influenced our results. Using permutation analysis, we determined that the t-statistic associated with the positive regression slope surpassed the null distribution representing regression to the mean for tcVNS (p=0.0040) but not taVNS (p=0.054). These results indicate that tcVNS evoked larger improvements for individuals with lower-than-average sensory performance.

**Figure 3.**
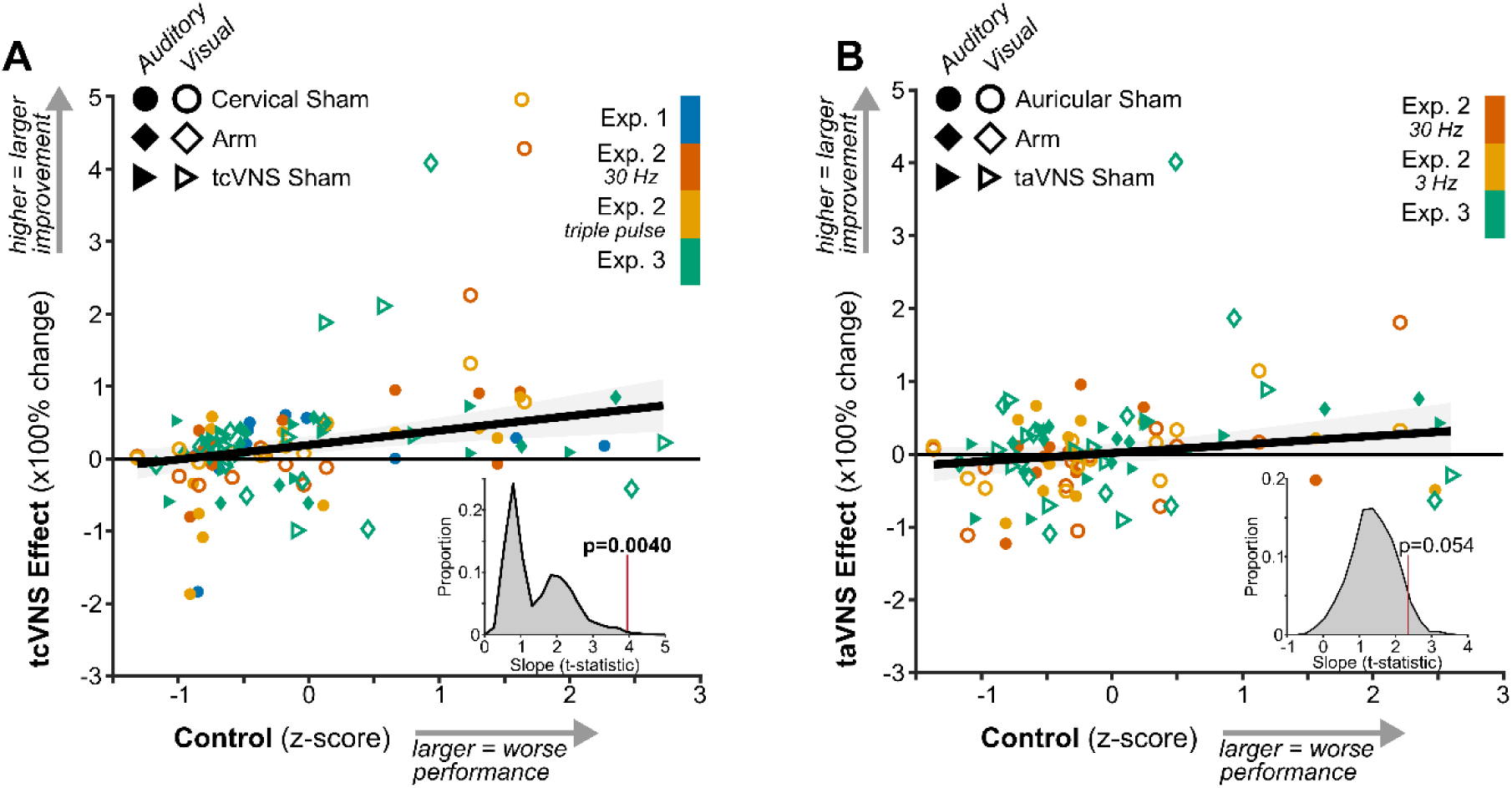
tcVNS-evoked improvements scaled with sensory performance. **(A)** Relation between sensory performance during control conditions (Cervical Sham, Arm, and tcVNS Sham) and the tcVNS effect – the percent change in sensory performance measured during tcVNS relative to the corresponding control condition. Each point depicts an individual participant in each experiment, sensory modality, and stimulation condition. The solid line depicts the best-fitting robust linear regression and the shaded region depicts its 95% confidence interval. The inset at bottom-right depicts the null distribution of t-statistics for the slope produced by regression to the mean. The vertical red line indicates the observed (i.e., non-permuted) t-statistic for the regression line and its associated p-value. **(B)** Follows the same conventions as A but depicts data for auricular tVNS.

### taVNS did not significantly improve sensory performance

taVNS did not evoke statistically significant improvements to auditory performance across two experiments (n=96 observations; p=0.67, Figure 4A; Table S5). Within Experiment 2, neither 30 Hz (t(11)=0.69, p=0.68, d [95% CI] = 0.20 [-0.44, 1.28]) nor 3 Hz taVNS significantly improved performance relative to Auricular Sham (t(11)=0.78, p=0.60, d [95% CI] = 0.23 [-0.16, 0.62]; Figure 4B). Likewise, within Experiment 3, performance during taVNS 30 Hz did not differ significantly between Arm (t(11)=2.21, p=0.054, d [95% CI] = 0.64 [0.27 1.16]) or taVNS Sham (t(11)=0.52, p=0.66, d [95% CI] = 0.15 [-1.00, 0.42]; Figure 4C).

**Figure 4.**
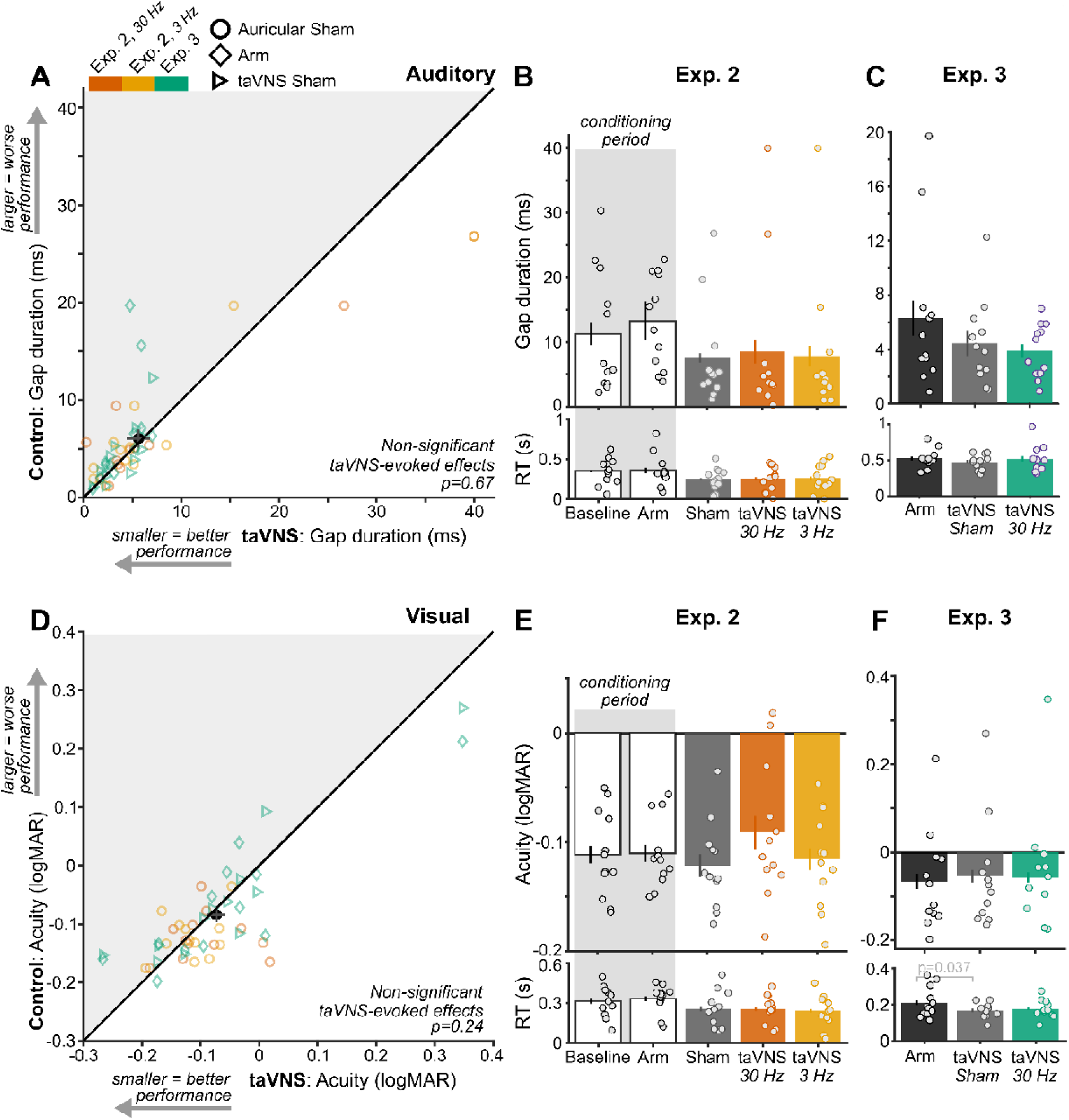
taVNS did not significantly improve sensory performance. **(A)** Gap duration thresholds during control conditions (Auricular Sham, Arm, and taVNS Sham) and active taVNS conditions across all experiments and stimulation protocols (30 Hz and 3 Hz). Plotting conventions follow those in Figure 2A. **(B)** Gap duration thresholds in each stimulation condition of Experiment 2 and **(C)** Experiment 3. Plotting conventions follow those in Figure 2B. **(D)** Visual acuity thresholds during control conditions and taVNS conditions across experiments and stimulation protocols. Plotting conventions follow those in A. **(E)** Letter size thresholds in each stimulation condition of Experiment 2 and **(F)** Experiment 3.

Similarly, taVNS did not generate statistically significant study-wide visual improvements (p=0.24, Table S6; Figure 4D). Neither taVNS 30 Hz (t(11) =-1.45, p=0.19, d [95% CI] =-0.42 [-0.94, 0.16]) nor taVNS 3 Hz significantly changed visual acuity relative to Auricular Sham in Experiment 2 (t(11)=0.43, p=0.69, d [95% CI] = 0.12 [0.028, 0.28]; Figure 4E). In Experiment 3, taVNS 30 Hz did not significantly alter visual acuity relative to Arm (t(11)=0.42, p=0.70, d [95% CI] = 0.12 [-0.54, 0.73]) or taVNS Sham (t(11)=0.16, p=0.89, d [95% CI] = 0.047 [-0.64, 0.65]). Due to the lack of significant taVNS-evoked performance changes, speed-accuracy tradeoffs were not assessed. That being said, RT was significantly faster during taVNS Sham than Arm stimulation in the visual task of Experiment 3 (t(11)=2.44, p=0.037; Figure 4F).

We further analyzed how effect sizes observed for taVNS compared to the corresponding value in our a priori power analysis (d=0.77) and to those observed for tcVNS. Effect sizes, pooled across experiments and sensory tasks, enabled a study-wide comparison between cervical and auricular tVNS efficacy. The effect size and confidence interval for taVNS (median=0.10, 95% CI = [-0.69 0.61]) were below the effect size in the power analysis. In contrast, the confidence interval for tcVNS effect size overlapped the value in our power analysis (median=0.54, 95% CI = [0.17 1.04]). Therefore, unlike tcVNS, taVNS did not elicit statistically significant improvements on sensory performance in our study.

### Behavioral state-dependent tVNS effects on heart rate variability

HRV, a cardiac indicator of vagal engagement, increased during tVNS relative to control stimulation, but only during passive behavioral states (Figure 5). Given that high cognitive demands, including performing sensory tasks, reduces HRV^38–41^ we measured HRV during two behavioral states – active task engagement and passive viewing. Interbeat intervals oscillated throughout each state (Figure 5A) and total spectral power at low (LF) and high frequencies (HF) indexed modulations of vagal activity^42–44^ (Figure 5B).

**Figure 5.**
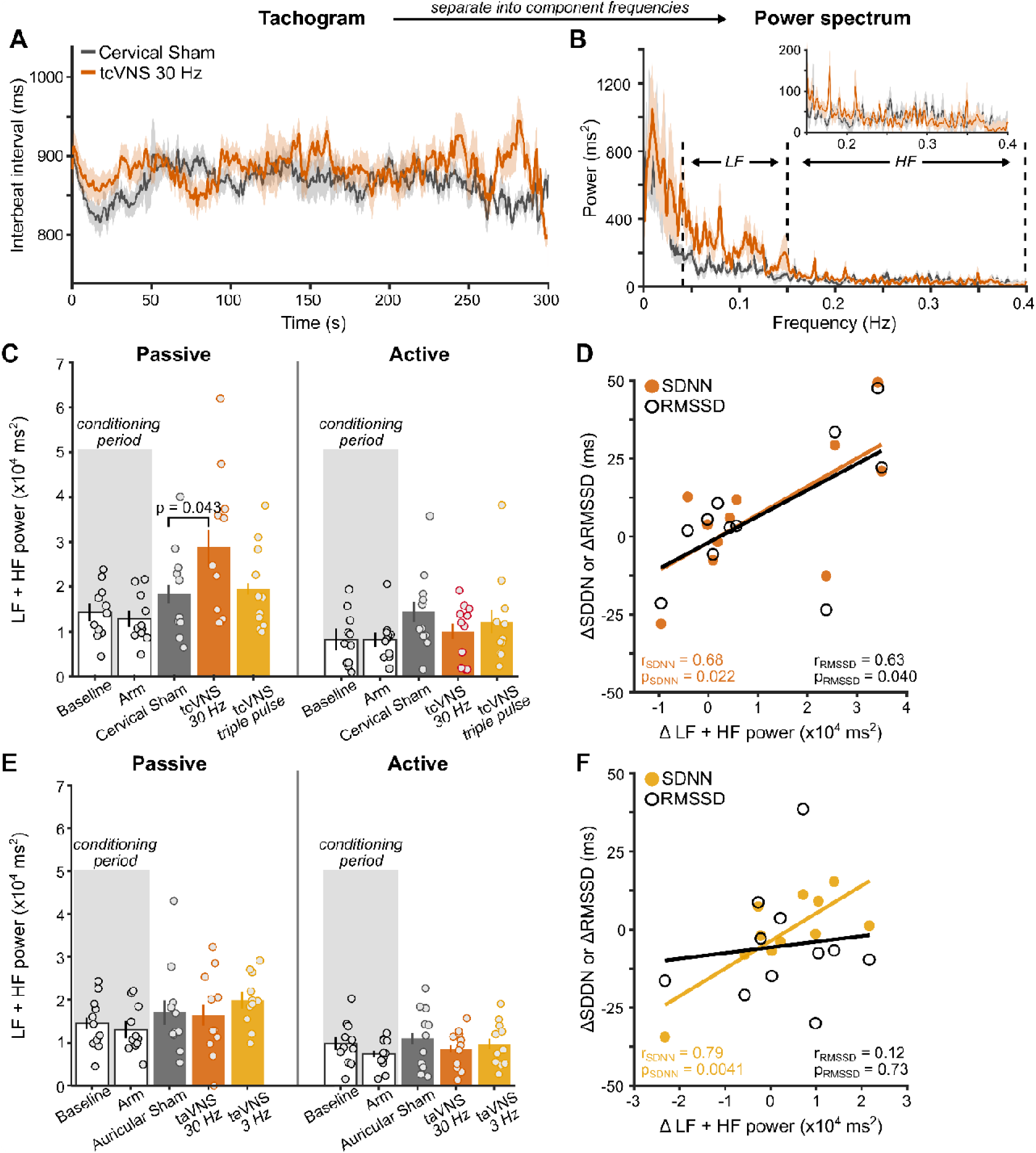
Behavioral state-dependent tVNS effects on heart rate variability. **(A)** Interbeat interval tachogram during passive viewing of Experiment 2. Solid lines show group-average and shaded regions depict ±1 within-subject SEM^34^. **(B)** Group-average power spectrum of the tachogram. Dashed lines demarcate boundaries for low (LF) and high frequency (HF) ranges. The inset shows a zoomed-in view of the HF range. **(C)** Summed LF and HF power for tcVNS during passive and active behavioral states. P-values denote results of permutation paired t-tests among sites. **(D)** Correlation among spectral and temporal HRV metrics. Points depict the difference between Cervical Sham and tcVNS 30 Hz in the frequency and temporal domain. Solid lines represent best-fitting linear regression to the data. **(E)** Summed LF and HF power for taVNS. **(F)** Correlation among spectral and temporal HRV metrics. Points depict the difference between Auricular Sham and taVNS 3 Hz.

Spectral power was significantly higher when participants exerted less cognitive effort during passive viewing^38–41^ (Figure 5C; F(1, 10)=16.582, p=0.0022, η^2^_G_=0.62). Differences among sites interacted with behavioral state (F(2, 20)=6.51, p=0.014, η^2^_G_=0.39). tcVNS did not alter HRV during active task engagement. During passive viewing, tcVNS 30 Hz significantly increased power relative to Cervical Sham (t(10)=2.48, p=0.043, d [95% CI] = 0.72 [0.045, 0.82]). Temporal domain metrics (SDNN, RMSSD) that similarly reflect modulations of vagal activity corroborated tcVNS-evoked increases in HRV (Figure 5D). Specifically, the difference between Cervical Sham and tcVNS 30 Hz in each HRV metric (total power, SDNN, and RMSSD) indexed stimulation-evoked effects. Increases in power were significantly correlated with increases in SDNN (r(9)=0.68, p=0.022, 95% CI = [0.13 0.91]) and RMSSD (r(9)=0.63, p=0.040, 95% CI = [0.041 0.89]). These findings suggest that HRV during passive states provided a clearer indication of vagal engagement by tcVNS.

taVNS did not generate state-dependent HRV changes (Figure 5E). Although spectral power was significantly higher during passive than active behavioral states (F(1, 10)=18.54, p=0.0015, η^2^_G_=0.65), neither taVNS 30 Hz (p=0.99) nor taVNS 3 Hz increased HRV relative to Auricular Sham during passive viewing (p=0.23). Likewise, temporal domain metrics did not reliably reflect taVNS-evoked modulations of vagal activity (Figure 5F). Here, we used differences between Auricular Sham and taVNS 3 Hz to index stimulation-evoked changes because taVNS 3 Hz evoked a small numerical increase in HRV. Although increases in power correlated significantly with increases in SDNN (r(9)=0.79, p=0.0041, 95% CI = [0.35 0.94]), it did not correlate with RMSSD (r(9)=0.12, p=0.073, 95% CI = [-0.52 0.67]). Thus, only tcVNS reliably increased HRV during a passive state.

### taVNS elicited discomfort in participants at levels initially comfortable

After each testing block, participants rated the discomfort experienced during stimulation (Figure 6A). Control conditions (Baseline, Arm, tcVNS Sham, and taVNS Sham) produced little to no discomfort in Experiments 2 and 3 (Figure 6B-C). Similarly, cervical stimulation conditions (Cervical Sham, tcVNS 30 Hz, and tcVNS triple pulse) produced only slight discomfort in a maximum of 7% of participants (Figure 6D). Conversely, auricular stimulation conditions produced moderate or higher levels of discomfort in up to 25% of participants in Experiments 2 and 3 (Figures 6E-F). Importantly, we followed the same procedue to identify the maximum intensity deemed comfortable in each condition, which was kept the same throughout each testing block.

**Figure 6.**
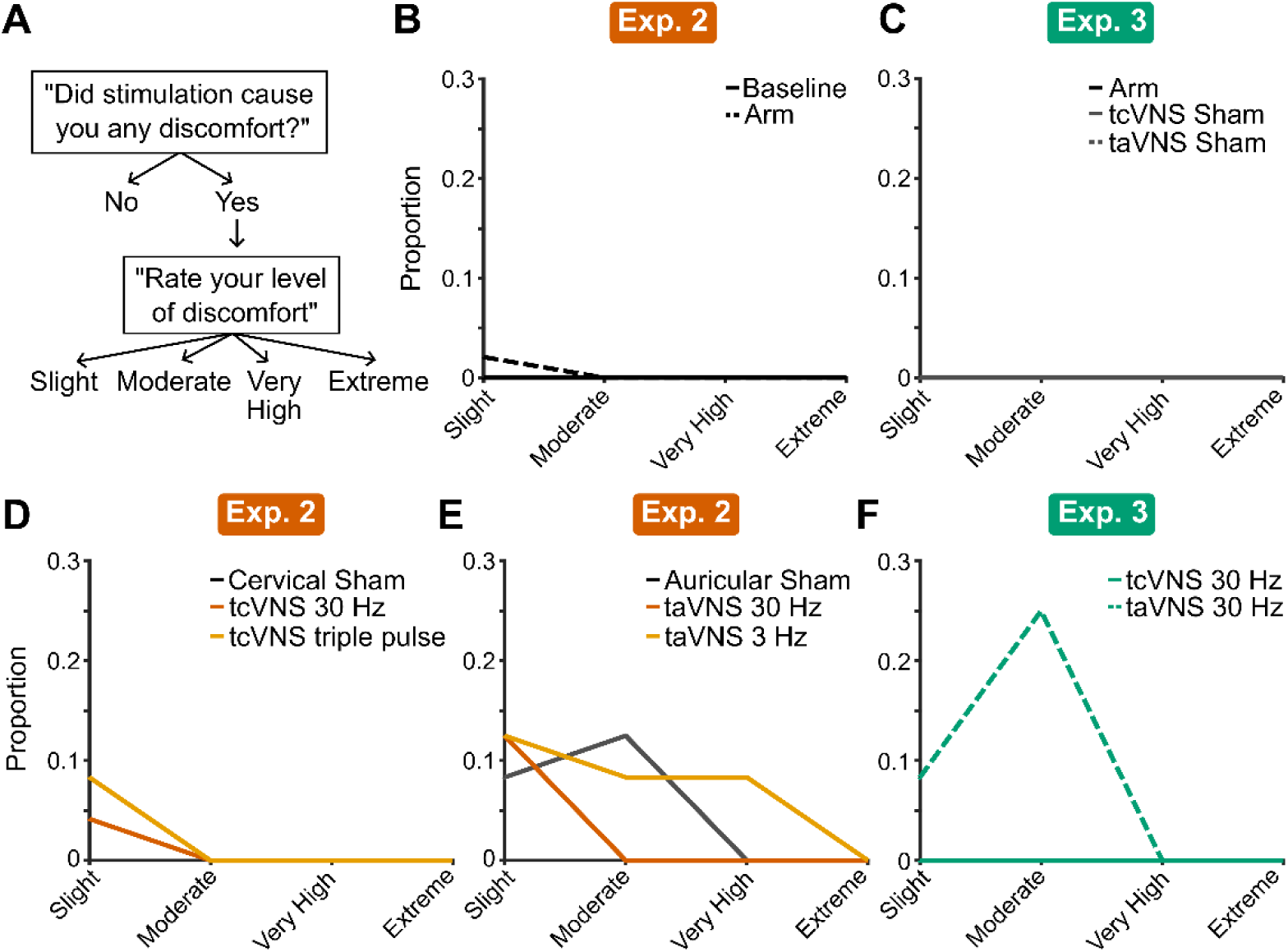
Participant discomfort potentially confounded taVNS effects on sensory performance. **(A)** Flowchart of the survey question and corresponding response options. **(B-F)** Histograms of participant responses for each experiment and stimulation protocol.

### Practice effects did not impact tVNS-evoked sensory improvements

We ruled out that repeatedly performing the auditory (Figure 7A-C) and visual tasks (Figure 7D-F) improved sensory performance, independent from tVNS effects. Sensory thresholds were compared across testing blocks when arranged in chronological order. Auditory gap duration thresholds decreased during the conditioning period of Experiment 2 (F(3, 33)=6.75, p=0.0049, η^2^_G_=0.38; Figure 7A). Relative to the first conditioning block, gap duration thresholds were significantly lower on the third (t(11)=2.99, p=0.017) and fourth blocks (t(11)=3.73, p=0.0037). However, thresholds did not decrease significantly after the second conditioning block, indicating that participants only required a single conditioning block to become proficient with the auditory task. By excluding the conditioning period from further analyses, we minimized contributions of practice effects to our results. Gap duration thresholds obtained after the conditioning period were stable across repeated testing blocks (Figure 7B). Participants in Experiment 3 completed a single training block prior to the main experiment, which effectively removed practice effects in the auditory task (Figure 7C).

**Figure 7.**
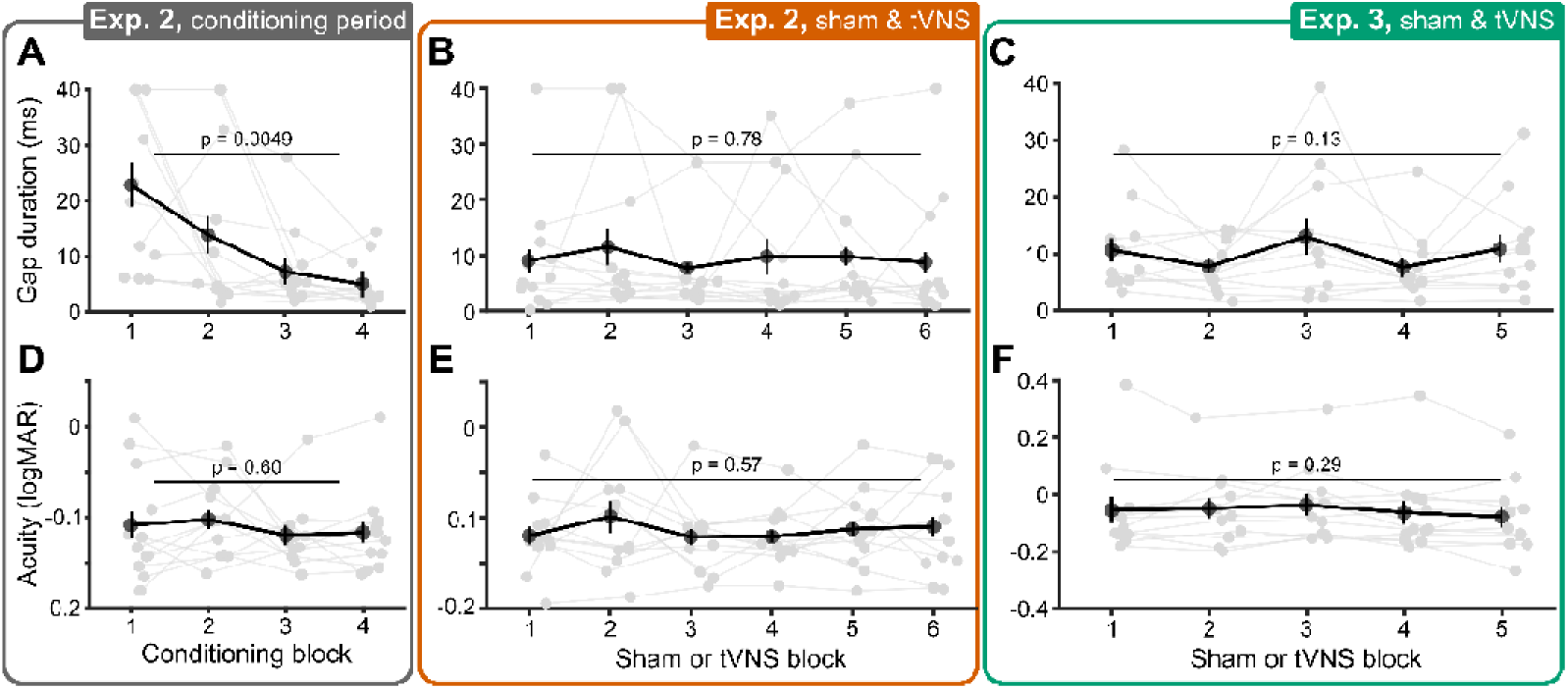
Practice effects did not impact tVNS-evoked sensory improvements. **(A)** Auditory gap duration thresholds ordered in chronological order of conditioning blocks in Experiment 2. Gray dots depict individual participants, black dots and error bars denote the group-average and ±1 within-subject SEM^34^, respectively. P-values denote the result of repeated measures ANOVAs. **(B)** Same as A, but showing thresholds during Sham and tVNS blocks, ordered chronologically in Experiment 2. **(C)** Same as B, but for Experiment 3. **(D)** Visual acuity thresholds ordered in chronological order of conditioning blocks in Experiment 2. **(E)** Same as D, but showing thresholds during Sham and tVNS blocks. **(F)** Same as E, but for Experiment 3.

The visual task was not impacted by practice, even during the conditioning period of Experiment 2 (Figure 7D-F). Participants were immediately proficient with the visual task, potentially indicating that it was more familiar and easier to learn. We additionally ruled out that the randomization procedure used in Experiments 2 and 3 biased our results toward observing tVNS effects (see “Supplementary Analyses”).

## Discussion

We measured the effects of continuous tVNS on sensory performance in neurotypical adults. tcVNS improved auditory gap discrimination by 37% and visual letter discrimination by 23%, on average. Individuals with lower sensory performance during control conditions experienced larger improvements during tcVNS. Conversely, taVNS did not significantly improve sensory performance. Lastly, tcVNS increased HRV relative to sham stimulation during passive viewing, corroborating vagal engagement. These results demonstrate that continuous tcVNS can enhance sensory performance in humans, particularly those with relatively poor sensory capabilities.

Our study design facilitated a comparison of the effectiveness of tcVNS and taVNS in evoking sensory improvements. While tcVNS significantly improved sensory performance and evoked the most substantial improvements for individuals with the lowest baseline sensory performance, taVNS did not generate statistically significant effects in auditory and visual performance. Moreover, tcVNS effect sizes aligned with our a priori power analysis that was based on results in animal models, suggesting that tcVNS is similarly effective at improving sensory performance as invasive stimulation in rodents. Conversely, taVNS effect sizes were negligible compared to our power analysis. These null findings add to an existing body of evidence showing mixed findings of the ability of taVNS to reliably evoke effects on cognitive performance and vagal activation biomarkers^28,45^. In addition, the lack of taVNS effects on HRV in our study aligns with recent meta-analysis findings^46^ that report no support of taVNS affecting vagally-mediated HRV. We also found our study design resulted in taVNS becoming uncomfortable during the trial for many participants, which did not occur with tcVNS. One potential is that the distraction of this pain could have obfuscated effects of taVNS on sensory processing. Overall, our results highlight the effectiveness of tcVNS in evoking sensory improvements and modulating HRV.

Our findings substantiate foundational studies in rodents. Steady activation of the LC-NE system through continuous LC stimulation^4^ or indirectly via continuous invasive VNS^5^ produced rapid and sustained improvements in sensory processing without affecting neural or behavioral response latencies. Improvements were driven by NE-mediated suppression of calcium T-type channels responsible for bursting activity by sensory relay neurons in the thalamus that reduces the accuracy and efficiency of sensory transmission^4–6^. Here, we show that tcVNS can induce similar perceptual performance enhancements in humans, suggesting that peripheral vagus nerve activation could induce NE-mediated sensory improvements in both species.

Notably, in both rodents^4,5^ and in the present study in humans, continuous VNS generated rapid and steady sensory benefits during stimulation delivery that generalized across different experiments, sensory tasks, and stimulus properties. In the current study, we randomized stimuli across trials (i.e., different letters and tone frequencies in the visual and auditory tasks, respectively) to minimize learning, and still observed tVNS-evoked improvements. These findings contrast with phasic VNS protocols common in motor and sensory learning protocols that repeatedly pair specific (i.e., non-randomized) stimuli with short VNS bursts over several days or weeks and induce relatively delayed and lasting sensory improvements specific to the paired stimulus^47–57^. Thus, our findings position continuous tcVNS as a new research avenue that can lead to tools for promoting general and on-demand improvements in sensory processing.

tcVNS-evoked increases in HRV provided evidence that our stimulation protocol engaged the vagus nerve^29,42–44^. We used common HRV indices, namely the power within low (LF) and high frequencies (HF), RMSSD, and SDDN. Although LF power has been historically associated with cardiac sympathetic activity, rather than vagally-mediated parasympathetic activity, mounting evidence indicates that the LF component is positively associated with vagal control^42–44^. Additionally, although SDNN is a common HRV measure that reflects mostly vagal activity^28,29^, it can also partly represent sympathetic cardiac activity.

Cardiac activity is known to be regulated by the vagus nerve^58^ and invasive stimulation of the vagus nerve can modulate HRV^59^. However, reports of tVNS effects on cardiac activity show conflicting results^28,60,61^. In this study, we demonstrated that tcVNS effects on HRV depended on behavioral state – tcVNS increased HRV during passive, but not active states. Consistent with earlier work^38–41^, heightened cognitive demands during task engagement diminished HRV, which potentially reduced the likelihood of observing tVNS-evoked effects. Thus, behavioral state exerts a strong influence on HRV and should be considered when measuring tVNS-related effects, alongside prior recommendations^28^. Critically, the observed tcVNS-evoked behavioral and cardiac effects correspond with animal work that have established their correlation with non-invasive readouts of LC-NE activity (i.e., pupil size)^4,6,62–65^. Therefore, our findings suggest that tcVNS altered vagal activity and subsequently modulated brain function, via the LC-NE, to improve human sensory processing.

Although it is well-established that invasive VNS subsequently activates the LC-NE system^30,66,67^, evidence that tVNS in humans can also activate the LC-NE system is mixed. Using functional magnetic resonance imaging, tVNS has been shown to evoke activity consistent with the location of the LC^21,23^ and in the nucleus tractus solitarius, which projects to the LC^22^. However, these studies did not use methods that appropriately delineate the LC^68,69^. Moreover, a comprehensive review^28^ of studies measuring non-invasive NE biomarkers (pupil diameter, P300, and salivary alpha amylase) found inconclusive evidence. Although we observed tcVNS-evoked sensory improvements consistent with findings from direct LC activation in rodents^4^, our study did not examine biomarkers of LC-NE activity (e.g., pupil size). Therefore, further work is required to confirm that the LC-NE system played a central role.

To strengthen our findings’ validity, several controls were assessed. Within individual experiments, tcVNS-evoked improvements were significant relative to two controls – stimulation to the upper trapezius (Cervical Sham) or withholding current from electrodes placed on the tcVNS site (tcVNS Sham). Although the arm stimulation control produced similar levels of sensory performance, it was more variable across individuals, which resulted in non-significant results. Larger variability necessitates larger sample sizes to observe significant tcVNS-evoked improvements relative to these controls. Therefore, the reliability of Cervical Sham and tcVNS Sham position them as effective controls for assessing tcVNS-evoked effects, particularly in small samples.

Previous tVNS studies in humans used a variety of stimulation parameters to assess tcVNS and taVNS effects^26^. tcVNS studies on cognitive performance^70,71^ and physiological markers^72–74^ typically delivered a sequence five sinusoidal pulses (pulse width: 200 μs/phase; 1 ms total) at 25 Hz to the cervical branch of the vagus nerve for up to 5 minutes. In the present study, we delivered either single biphasic pulses (pulse width: 200 μs) or a sequence of three biphasic pulses (pulse width: 600 μs) at 30 Hz. Critically, we delivered stimulation continuously throughout each 10-15 minute long testing block to mimic the continuous LC-NE activation shown to underlie sensory improvements in animal models^4–6^. Therefore, by using stimulation parameters based on findings in animal models, we observed similar sensory improvements in humans.

Our taVNS stimulation protocol was unlike previous taVNS studies that utilized electrodes placed on the ear’s lateral surface^26^. Our approach involved placing electrodes on both the lateral and medial ear surfaces. We hypothesized that this configuration would enhance current flow through the tissue and better activate the vagus nerve. Moreover, our Auricular Sham site also differed from the commonly used earlobe site^26^ as it has been suggested the earlobe is not a good taVNS sham^75^. Therefore, it remains an open question whether electrode placement on only the lateral surface and a control site on the earlobe may have revealed taVNS-evoked improvements while also reducing participant discomfort. Future investigations could shed light on the impact of taVNS electrode placement and vagus nerve mediated neuromodulation.

Surprisingly, our taVNS protocol elicited moderate or very high discomfort in 25% of participants whereas tcVNS produced a comfortable experience with no moderate or high discomfort reported. To note, both conditions followed the same procedure to determine a max comfortable level for each participant that was then fixed for the duration of that session. The experienced discomfort could have potentially influenced taVNS efficacy. Beyond being distracting from performing the sensory tasks, discomfort and pain are known to decrease vagal activity^76^, which may have diminished the ability of taVNS to modulate sensory performance and weakened taVNS-evoked effects on HRV. Additional research is needed to delineate the interaction between comfort and tVNS-evoked effects and our findings demonstrate the utility of considering individuals’ subjective experience during tVNS.

Lastly, we explored the effectiveness of two tcVNS (30 Hz and triple pulse) and taVNS protocols (30 Hz and 3 Hz) guided by prior studies with animal models demonstrating they modulated LC activity^30^. Both taVNS protocols did not evoke statistically significant behavioral and HRV effects whereas tcVNS protocols yielded numerically similar but statistically different behavioral and HRV effects. tcVNS 30 Hz significantly increased HRV, auditory, and visual performance relative to controls whereas tcVNS triple pulse significantly improved visual performance relative to a subset of controls. The sample size within each experiment was determined via a priori power analyses and may have been insufficient to measure statistically significant differences among all stimulation protocols. Therefore, it is possible that each tcVNS protocol yields similar statistical results at larger sample sizes. Previous work with finite element modeling provided insights regarding the effectiveness of common tcVNS protocols in activating fiber subtypes of the vagus nerve^77^. Similar computational modeling work could further shed light on whether each waveform (single pulse vs triple pulse) differentially drives vagal activity.

In conclusion, we demonstrated that continuous tcVNS can rapidly enhance sensory performance in humans. These findings provide evidence that neuromodulation via tcVNS may be a useful tool for immediately augmenting hearing and vision, particularly for individuals with relatively impaired sensory capabilities. Moreover, these results suggest more extensive testing is warranted to assess the clinical relevance of using tcVNS to induce NE-mediated enhancement of central sensory processing. Overall, this study bridges the gap between animal and human research and offers promising implications for the development of new therapies for sensory disorders.

## Methods

### Ethics

Participants were passively recruited, provided written informed consent, and compensated $30/hour. All experimental protocols complied with the Declaration of Helsinki and were approved by Cornell University’s Institutional Review Board for Human Participant Research. Informed consent was provided for the images in Figure 1A to be published in an online open-access publication.

### Sample

Twenty-nine individuals participated across three experiments, with 12 participants per experiment (18 females; age: 28±4.9 (SD); range: 18-39). Most participants completed a single experiment (ten in Exp.1, seven in Exp.2, seven in Exp.3), three completed Experiments 2-3, and two completed all three experiments. Participants were asked to refrain from any stimulants (e.g., caffeine, nicotine) for three hours prior to participating and were naïve to the study’s purpose and the location of the vagus. All participants passed inclusion criteria for the study: no substance abuse history, no neurological or psychiatric disorders, no cardiac disorders, no medical implants, no prior abnormalities or surgeries involving the neck or vagus nerve, and normal or corrected-to-normal hearing and vision. The sample size for each individual experiment was determined a priori using G*Power 3.1.9.4^78^ based on a published study^4^. The study discovered the mechanism of action underlying effects of continuous LC-NE activation on sensory performance in rodents. It employed a stimulation pattern and sensory discrimination task similar to those in our current study, was corroborated by further invasive VNS work in rodents^5^, and was agnostic regarding the vagal branch being stimulated, making it a preferable basis for our power analysis. Specifically, the study demonstrated an effect size (Cohen’s d^79,80^=0.77) estimated from a paired t-test comparing behavioral performance of rodents performing a tactile discrimination task during LC stimulation versus sham. We were interested in detecting similarly large effect sizes for tcVNS and taVNS. Assuming tVNS drives LC-NE activity to improve human sensory performance with a similar effect size, α=0.05, and power=0.8, 12 would be the minimum sample size required. To strengthen the validity of our findings, we pooled data from all three experiments for our critical analyses (see “Sensory Performance”), utilizing our complete sample of 29 participants and thereby increasing statistical power.

### Stimulation

A Mettler Sys*Stim 240 neuromuscular stimulator (Mettler Electronics Corporation, CA, USA) set to constant current mode delivered current through a pair of gel electrodes (diameter: 1”; PALS, Axelgaard Manufacturing, CA, USA) placed at one of 5 sites (Figure 1A). We delivered two waveforms, a single biphasic square pulse (100 μs/phase; 200 μs total) delivered continuously at one of two frequencies (30 Hz, 3 Hz), or a sequence of three biphasic square pulses (100 μs/phase; 600 μs total) delivered at 30 Hz (Figure 1B). To determine current amplitude (Table 1), current intensity was gradually ramped up until participants reported muscle contractions or pain, after which it was reduced to the maximum level that was perceptible, tolerable, and did not induce muscle activation. Prior to delivering stimulation, sites were cleaned with a hypoallergenic sanitary wipe.

**Table 1.** Current amplitude delivered to each stimulation site in each experiment.

| Experiment | Site | Current (M ± SD) |
| --- | --- | --- |
| 1 | Cervical Sham | 7.8 ± 1.7 mA |
| | tcVNS 30 Hz | 6.5 ± 1.5 mA |
| 2 | Arm | 5.1 ± 1.6 mA |
| | Cervical Sham | 5.4 ± 2.0 mA |
| | Auricular Sham | 2.5 ± 2.0 mA |
| | tcVNS 30 Hz | 5.5 ± 1.6 mA |
| | tcVNS Triple pulse | 5.1 ± 1.9 mA |
| | taVNS 30 Hz | 2.3 ± 1.6 mA |
| | taVNS 3 Hz | 2.5 ± 1.8 mA |
| 3 | Arm | 6.9 ± 1.2 mA |
| | tcVNS 30 Hz | 7.1 ± 1.2 mA |
| | taVNS 30 Hz | 2.0 ± 0.9 mA |

Ten stimulation conditions were assessed:

#### Active

1. **tcVNS 30 Hz**. Electrodes were placed within the left carotid triangle, lateral to the larynx, medial to the sternocleidomastoid muscle (SCM), and oriented parallel to the SCM (3-5 cm center-to-center) to target the cervical branch of the vagus nerve^8,9^. Pulses were delivered at 30 Hz.
2. **tcVNS Triple pulse**. Electrodes were positioned identically to tcVNS 30 Hz, and waveforms comprising three biphasic pulses were delivered at 30 Hz.
3. **taVNS 30 Hz**. Electrodes were placed on the lateral and medial surfaces of the left ear to direct current flow through the ear and into the auricular branch of the vagus nerve (ABVN). Based on cadaveric dissections^7^, one electrode was placed on the lateral surface overlapping the cymba concha and cavum concha, and the second was placed on the middle-third of the dorsal medial surface. Pulses were delivered at 30 Hz.
4. **taVNS 3 Hz**. Electrodes were positioned identically to taVNS 30 Hz, but pulses were delivered at 3 Hz.

#### Control

1. **Baseline**. No electrodes were placed on the participant; accordingly, no stimulation was delivered.
2. **Arm**. Electrodes were positioned on the left forearm (3-4 cm center-to-center), approximately over the brachioradialis muscle. Pulses were delivered at 30 Hz.
3. **Cervical Sham**. Electrodes were placed near the junction between the rear of the neck and the shoulder (3-4 cm center-to-center), overlapping the trapezius muscle, which is primarily innervated by efferent fibers of cranial nerves II, III, IV, and XI, but not by the vagus nerve^81^. Pulses were delivered at 30 Hz. Because stimulation of cranial nerve II afferents can improve cognition^82^, any tVNS-mediated improvements we observe relative to this sham site are potentially underestimated.
4. **Auricular Sham**. Electrodes were placed on the superior crus of antihelix, with one electrode on the ear’s lateral surface, and another on the medial surface. The great auricular nerve innervates the lateral location and the ABVN does not run through the medial location in cadaveric dissections^7^. Pulses were delivered at 30 Hz.
5. **tcVNS Sham**. Electrodes were positioned identically to tcVNS 30 Hz, but current was not delivered. To maintain participant blinding, current was initially ramped up to a low but perceptible level for 30-60 seconds, then ramped down to 0 mA prior to the start of sensory tests.
6. **taVNS Sham**. Electrodes were positioned identically to taVNS 30 Hz and followed the ramp-up, ramp-down procedure of tcVNS no-stim.

### Procedure

Stimulation was delivered continuously throughout each testing block, except during controls without stimulation (Baseline, tcVNS Sham, taVNS Sham). Across experiments, participants completed auditory (Experiments 1-3) and visual tasks (Experiments 2-3; Figure 1C), or passively viewed a blank computer display with no associated task for five minutes (Experiment 2). This passive viewing period enabled assessments of HRV while participants had low cognitive demands; heightened demands during active task engagement are known to decrease HRV^38–41^ and may obscure tVNS effects on HRV. During sensory tasks, adaptive procedures determined auditory and visual thresholds that indexed sensory performance (Figure 1D). Then, participants completed brief surveys (Experiments 2-3).

All experiments followed crossover, sham-controlled, and single-blind designs, but differed in the number of stimulation and sham conditions (Figure 1E). Each stimulation condition lasted 15-20 minutes with stimulation being delivered continuously throughout, and experiments were performed based on participant availability, typically in the morning (9:30 am – 12 pm) or afternoon (1 pm – 6 pm). Experiment 1 included Cervical Sham and tcVNS 30 Hz sessions with their order counterbalanced across participants, completed in a single day, and separated by a 15-30 minute washout period (M=20.23, SD=3.68 min). Experiment 2 included eight conditions, split into cervical and auricular sessions completed on separate days in a randomized order. Within each session, five conditions were completed in pseudo-randomized order separated by 10-25 minute washout periods (M=13.96, SD=4.09 min): Blocks 1 and 2 contained Baseline or Arm, in randomized order, to serve as a conditioning period that minimized apprehension toward the stimulation sensation and acclimated participants to the sensory tasks. Then in blocks 3-5 of cervical VNS sessions, Cervical Sham, tcVNS 30 Hz, and tcVNS triple pulse were completed in randomized order. Alternatively, during blocks 3-5 of auricular VNS sessions, Auricular Sham, taVNS 30 Hz, or taVNS 3 Hz were completed in randomized order. Experiment 3 began with a practice block, without stimulation, that familiarized participants with the sensory tasks. Then five conditions were completed in a single day in randomized order, separated by 15-40 minute washout periods (M=23.59, SD=10.28 min) – Arm, tcVNS 30 Hz, taVNS 30 Hz, tcVNS Sham, and taVNS Sham.

### Apparatus

Psychtoolbox-3^83^ generated sensory stimuli in MATLAB (MathWorks, MA, USA). An Empatica E4 wristband (Empatica Inc., MA, USA) equipped with photoplethysmography was positioned on participants’ left wrist during Experiment 2 to monitor HRV. All experiments were conducted with a Lenovo ThinkStation running Windows 10. A 24 inch (Experiment 1 – ASUS VG245H; 1920×1080; 75 Hz) or 27 inch LCD display (Experiments 2 and 3 – ASUS VG27BQ; 2560×1440; 144 Hz) was used to display visual stimuli. Experiments 2-3 required a higher resolution with increased pixel density and refresh rate to support the letter discrimination task. Gamma correction was performed with a Datcolor SpyderX Pro colorimeter and calibration routines implemented in DisplayCAL (ArgyllCMS, displaycal.net). Auditory stimuli were delivered via factory-calibrated ER-2 insert earphones (Etymotic Research Inc), connected to a digital-to-analogue converter (Atom DAC+, JDS Labs) and amplifier (Atom Amp+, JDS Labs). Participants sat in a chair positioned 80-90 cm from the display. Visual stimuli were scaled to maintain an identical visual angle at all viewing distances. Auditory volume was set manually by the experimenter to the highest level deemed comfortable by the participant at the beginning of each session and was not adjusted within sessions.

### Psychophysics

We used psychophysics to evaluate tVNS effects on sensory performance, guided by our prior animal research showing enhanced thalamic sensory processing during invasive VNS. We tested auditory and visual modalities as both are reliant on thalamic processing.

*Auditory*. Participants performed an auditory gap discrimination task commonly used to assess central auditory temporal processing^84–87^, and similar to a commercially available Random Gap Detection Test^88^. To control eye movements, participants maintained visual fixation on a dot (luminance: 0.23 cd/m^2^) presented on a mid-gray background (55 cd/m^2^). Participants reported which of two temporal intervals contained a tone with a gap. 500 ms of silence separated intervals. Auditory stimuli were pairs of 17 ms pure tones (Experiment 1 used 1000 or 2000 Hz in separate blocks; Experiments 2-3 used 10 frequencies logarithmically spaced between 1000-4000 Hz, interleaved within blocks). One tone-pair had no gap while another was separated by silence. The tone-pair containing a gap was randomly determined on each trial. Participants reported which tone-pair contained a gap. Inter-trial intervals followed responses (3000 ms for Experiment 1; 500 ms for Experiments 2-3).

*Visual*. Participants performed a letter discrimination task adapted from clinical measures of visual acuity – Bailey-Lovie and ETDRS charts^36^. Participants maintained visual fixation on a central dot (0.23 cd/m^2^) presented on a white background (228 cd/m^2^) and reported which letter appeared on-screen for 500 ms. Letters were displayed in Sloan font, as used in commercially available eye charts^36^. Participants responded by pressing the corresponding letter on a keyboard after stimulus offset. One of ten optotypes appeared on each trial – C, D, H, K, N, O, R, S, V, Z. Participants were not informed that only these letters would appear and could respond with any letter of the alphabet. A 500 ms inter-trial interval followed responses.

For all sensory tasks, participants were instructed to respond with their initial reaction, within a 3-second time window after stimulus offset. Additionally, participants first completed practice trials during which feedback was provided. During practice, participants received feedback on their performance to ensure they were able to perform the task correctly. Participants were required to complete these relatively easy practice trials until they did three in a row without error. In Experiment 3, participants underwent a practice block comprising 15 practice trials spanning all difficulty levels, without stimulation, while receiving trial-wise feedback to acquaint themselves with the task. Then, they completed 30 (Experiment 1) or 100 trials (Experiments 2-3) per stimulation condition without feedback during which task difficulty was adaptively controlled (see “Thresholding”). This procedure of providing then withholding feedback follows prior studies^89^.

### Thresholding

We used QUEST^90^, an adaptive procedure implemented in Palamedes toolbox^91^ and commonly used with psychophysics^92–96^ to adjust task difficulty based on participants’ response accuracy. Indices of task difficulty were gap duration and letter size for auditory and visual tasks, respectively. Generally, these procedures presented harder-to-discriminate stimuli after correct responses and easier-to-discriminate stimuli after incorrect responses, until they converged on a threshold stimulus that maintained 75% accuracy (Figure 1D). Letter size thresholds, in minutes of arc, from the visual task were transformed to logMAR units using the formula^36^: logMAR = log_10_(size ÷ 5).

Each QUEST procedure began with a calibration sweep that presented 10 (Experiment 1) or 5 (Experiment 2) pre-determined stimulus levels. These levels spanned a large range of possible stimuli and initialized each procedure near the participant’s threshold. In Experiment 1, a single QUEST procedure targeted sensory stimuli that would maintain 75% task accuracy (Figure 1D, top). For Experiments 2 and 3, five QUEST procedures targeted five linearly-spaced levels of accuracy ranging from near-chance (55% – auditory, 25% – visual) to near-perfect (95%). As a result, the presented stimuli sampled a large performance range. Logistic functions were fit to trial-wise sensory judgements and characterized participants’ psychometric functions – the mapping between stimulus (*x*) and response accuracy (*p*):

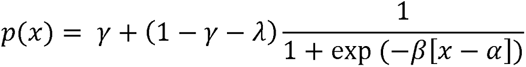

where γ is the lower asymptote set to chance-level performance determined by the task (50% in gap discrimination, 3.8% – in letter discrimination), λ is the upper asymptote determined by a participant’s maximum response accuracy, *β* is the function’s slope, and α is the midpoint between the asymptotes. Fits were optimized via maximum likelihood estimation – 100 trials constrained 3 free parameters. Thresholds for 75% accuracy were determined from the fitted functions (Figure 1D, bottom).

### Sensory performance

Our critical analyses comprised linear mixed models (LMM) and robust linear regression that pooled across experiments to examine study-wide trends whereas paired t-tests examined within-experiment comparisons. For LMMs and robust linear regression, sensory thresholds determined by QUEST procedures were merged across experiments. We additionally measured reaction time as a secondary dependent variable to assess speed-accuracy tradeoffs. Statistical analyses were conducted with the Statistics and Machine Learning toolbox in MATLAB R2021b.

LMMs evaluated whether tcVNS and taVNS improved auditory (cervical: n=120 observations; auricular: n=96) or visual task performance (n=96), relative to all control conditions except Baseline and Arm conditions that were completed within the conditioning period during Experiment 2 (Figure 1E). LMMs were configured with the formula:

Threshold ∼ 1 + Site + (1|Experiment) + (1|Participant)

where *Threshold* (dependent variable) referred to participants’ sensory thresholds. *Site* (fixed effect) was a categorical variable that indexed a condition as active or control. *Experiment* (random effect) classified each experiment; we additionally coded each active protocol (triple pulse, 30 Hz, 3 Hz) as separate experiments. Thus, *Experiment* accounted for variables outside of experimental control (e.g., stimulation sensation, time-of-day, demographics). *Participant* uniquely identified each participant, which preserved the within-subject design in cases where participants completed multiple experiments. LMM complexity was constrained by the available data and validated by positive definite Hessian matrices at model convergence. Due to a non-positive definite Hessian matrix at convergence, the (1|Experiment) term in the model formula was excluded for taVNS effects on visual acuity to maintain an appropriate level of model complexity for the available data (Table S6).

Robust linear regression evaluated whether sensory performance during control conditions, with the exception of Baseline and Arm conditions in Experiment 2, predicted the magnitude of tcVNS (n=108) or taVNS effects (n=96). A Cauchy weight function minimized outliers’ influence during regression. Data were merged across experiments and sensory tasks. We quantified the magnitude of tVNS-evoked effects as the proportion change in sensory thresholds between each active and control condition using the formula: tVNS effect = (control-tVNS) ÷ control. Sensory thresholds during control conditions were z-scored within each task, which put thresholds on a common scale and facilitated merging across tasks. To assess whether regression to the mean impacted our results, permutation analyses were conducted^97^. Labels were randomly permuted 1000 times among control and active conditions within participants, sensory tasks, stimulation sites, and experiments. Robust linear regression was performed on each permuted dataset and the resulting t-statistic of the regression coefficient populated a null distribution of regression to the mean. P-values were calculated as the number of t-statistics in the null distribution that exceeded that from the original, non-permuted, dataset.

Within each experiment, Shapiro-Wilk tests assessed normality. If normality was established, paired t-tests were used (Experiment 1). If not, permutation paired t-tests were used (100,000 permutations; Experiments 2-3). We corrected for multiple comparisons via the Max T method, which ensured strong control of the family wise error rate^98^. We utilized Cohen’s d^79,80^ for effect size quantification, reporting analytically-derived confidence intervals (Experiment 1) or bootstrapped 95% confidence intervals when permutation paired t-tests were conducted (Experiment 2-3) to assess compatibility with the effect size assumed in our power analysis.

### HRV

Interbeat intervals were measured during passive viewing and task engagement in Experiment 2. The passive viewing condition helped mitigate the negative impact of heightened cognitive demands on HRV^38–41^, preventing potential masking of tVNS-evoked HRV effects. Three HRV metrics were computed that each index modulations of vagal activity^29,42–44^: standard deviation of normal-to-normal intervals (SDNN), root mean square of successive differences between heartbeats (RMSSD), and summed spectral power within low (LF: 0.04-0.15 Hz) and high frequency (HF: 0.15-0.4 Hz) ranges. Repeated measures ANOVAs assessed the impact of *Site* and *State* (passive vs active), Greenhouse-Geisser corrected p-values are reported alongside the effect size quantified as generalized eta-squared (η^2^_G_)^80^. Permutation t-tests assessed differences among sites within each state and Cohen’s d quantified effect size. Pearson correlation quantified linear relations among HRV metrics.

### Survey

Participants completed surveys upon completing sensory tasks (Experiments 2-3). The survey asked “Did stimulation cause you any discomfort?” and provided two response options: “Yes” and “No”. If “Yes”, participants rated their discomfort using a Likert scale: 1=Slight, 2=Moderate, 3=Very High, and 4=Extreme.

### Practice Effects

To assess whether repeated execution of auditory and visual tasks improved sensory performance, independently from tVNS-evoked effects, sensory thresholds were compared across testing blocks when ordered chronologically. Repeated measures ANOVAs assessed the effect of *Testing Block* on sensory thresholds, separately for the auditory and visual tasks, and Greenhouse-Geisser corrected p-values are reported alongside generalized eta squared (η^2^_G_)^80^. For Experiment 2, sensory thresholds from the conditioning period (i.e., Baseline and Arm conditions) were assessed separately from Sham and tVNS conditions. In cases where a significant *Testing Block* effect was observed, permutation paired t-tests were performed between all pairs of testing blocks, p-values were corrected for multiple comparisons via the Max T method^98^.

## Abbreviations

LC: locus coeruleus
LC-NE: NE, locus coeruleus-norepinephrine system
tVNS: transcutaneous vagus nerve stimulation
taVNS: transcutaneous auricular vagus nerve stimulation
tcVNS: transcutaneous cervical vagus nerve stimulation
HRV: heart rate variability
SCM: sternocleidomastoid muscle
ABVN: auricular branch of the vagus nerve cd/m2, candelas per meter squared
LMM: linear mixed model
SDNN: standard deviation of normal-to-normal heartbeat intervals
RMSSD: root mean square of successive differences between heartbeat intervals
LF: low frequency
HF: high frequency
RT: reaction time

## Acknowledgements

The authors acknowledge the input of Dr. Frank Lin, M.D., Ph.D., on an earlier draft of this manuscript, Cynthia Poon for providing writing and proofreading assistance, and all individuals who participated in this study.

## CRediT authorship contribution statement

**M.J.:** Conceptualization, Methodology, Software, Validation, Formal analysis, Investigation, Data Curation, Visualization, Writing – Original Draft, Review & Editing,

**J.B.C:** Conceptualization, Supervision, Writing – Review & Editing

**Q.W.:** Conceptualization, Supervision, Writing – Review & Editing

**C.R.:** Conceptualization, Methodology, Supervision, Project administration, Resources, Funding acquisition, Writing – Review & Editing.

## Data availability

The datasets generated during and/or analyzed during the current study are available from the corresponding author, C.R., on reasonable request.

