## Supplementary Materials for "Transcutaneous cervical vagus nerve stimulation improves sensory performance in humans"

### Tables

**Table S1.** Summary table for the linear mixed model of the impact of tcVNS on auditory task performance.

##### Model Information

| | |
|---|---|
| Number of observations | 120 |
| Fixed effects coefficients | 2 |
| Random effects coefficients | 34 |
| Covariance parameters | 3 |

##### Formula

Threshold ~ 1 + Site + (1 | Participant) + (1 | Experiment)

##### Fixed Effects Coefficients

| Name | Estimate | SE | 95% CI | t-statistic | df | p-value |
|---|---|---|---|---|---|---|
| Control (intercept) | 10.17 | 1.86 | [6.49 13.85] | 5.47 | 118 | $2.51 \times 10^{-7}$ |
| Active (slope) | -3.75 | 1.05 | [-5.83 -1.67] | -3.57 | 118 | 0.00052 |

##### Random Effects Covariance Parameters

| Grouping Variable | Coefficient Names | Estimate | 95% CI |
|---|---|---|---|
| Participant (29 levels) | Control SD | 7.72 | [5.69 10.48] |
| Experiment (5 levels) | Control SD | 1.91 | [0.61 5.98] |
| Error | Residual SD | 5.76 | [4.96 6.68] |

**Table S2.** Summary table for the linear mixed model of the impact of tcVNS on visual task performance.

#### Model Information

| | |
|---|---|
| Number of observations | 96 |
| Fixed effects coefficients | 2 |
| Random effects coefficients | 23 |
| Covariance parameters | 3 |

#### Formula

Threshold ~ 1 + Site + (1 | Participant) + (1 | Experiment)

#### Fixed Effects Coefficients

| Name | Estimate | SE | 95% CI | t-statistic | df | p-value |
|---|---|---|---|---|---|---|
| Control (intercept) | -0.072 | 0.025 | [-0.12 -0.023] | -2.92 | 94 | 0.0044 |
| Active (slope) | -0.016 | 0.0077 | [-0.032 -0.00093] | -2.11 | 94 | 0.038 |

#### Random Effects Covariance Parameters

| Grouping Variable | Coefficient Names | Estimate | 95% CI |
|---|---|---|---|
| Participant (19 levels) | Control SD | 0.10 | [0.075 0.14] |
| Experiment (4 levels) | Control SD | 0.0041 | [0 0.36] |
| Error | Residual SD | 0.038 | [0.032 0.045] |

**Table S3.** Summary table for the robust linear regression relating the magnitude of tcVNS effects to sensory performance during control conditions.

#### Model Information

| | |
|---|---|
| Number of observations | 108 |
| Number of coefficients | 2 |

#### Formula

Effect ~ 1 + Control

#### Model Fit Statistics

| | |
|---|---|
| Root Mean Squared Error | 0.50 |
| F-statistic vs. constant model | 17.2 |
| P-value of F-statistic | $6.59 \times 10^{-5}$ |

#### Fixed Effects Coefficients

| Name | Estimate | SE | 95% CI | t-statistic | p-value |
|---|---|---|---|---|---|
| Intercept | 0.19 | 0.049 | [0.094 0.29] | 3.92 | $1.57 \times 10^{-4}$ |
| Control (slope) | 0.20 | 0.051 | [0.10 0.30] | 3.95 | $1.40 \times 10^{-4}$ |

**Table S4.** Summary table for the robust linear regression relating the magnitude of taVNS effects to sensory performance during control conditions.

#### Model Information

| | |
|---|---|
| Number of observations | 96 |
| Number of coefficients | 2 |

#### Formula

Effect ~ 1 + Control

#### Model Fit Statistics

| | |
|---|---|
| Root Mean Squared Error | 0.67 |
| F-statistic vs. constant model | 5.52 |
| P-value of F-statistic | 0.021 |

#### Fixed Effects Coefficients

| Name | Estimate | SE | 95% CI | t-statistic | p-value |
|---|---|---|---|---|---|
| Intercept | 0.049 | 0.068 | [-0.087 0.18] | 0.71 | 0.48 |
| Control (slope) | 0.17 | 0.072 | [0.026 0.31] | 2.35 | 0.021 |

**Table S5.** Summary table for the linear mixed model of the impact of taVNS on auditory task performance.

#### Model Information

| | |
|---|---|
| Number of observations | 96 |
| Fixed effects coefficients | 2 |
| Random effects coefficients | 23 |
| Covariance parameters | 3 |

#### Formula

Threshold ~ 1 + Site + (1 | Participant) + (1 | Experiment)

#### Fixed Effects Coefficients

| Name | Estimate | SE | 95% CI | t-statistic | df | p-value |
|---|---|---|---|---|---|---|
| Control (intercept) | 6.04 | 1.59 | [2.87 9.20] | 3.79 | 94 | 0.00027 |
| Active (slope) | -0.42 | 0.97 | [-2.35 1.51] | -0.43 | 94 | 0.67 |

#### Random Effects Covariance Parameters

| Grouping Variable | Coefficient Names | Estimate | 95% CI |
|---|---|---|---|
| Participant (19 levels) | Control SD | 5.08 | [3.50 7.39] |
| Experiment (4 levels) | Control SD | 1.66 | [0.50 5.54] |
| Error | Residual SD | 4.76 | [4.05 5.60] |

**Table S6.** Summary table for the linear mixed model of the impact of taVNS on visual task performance.

#### Model Information

| | |
|---|---|
| Number of observations | 96 |
| Fixed effects coefficients | 2 |
| Random effects coefficients | 19 |
| Covariance parameters | 2 |

#### Formula

Threshold ~ 1 + Site + (1 | Participant)

#### Fixed Effects Coefficients

| Name | Estimate | SE | 95% CI | t-statistic | df | p-value |
|---|---|---|---|---|---|---|
| Control (intercept) | -0.084 | 0.024 | [-0.13 -0.037] | -3.54 | 94 | 0.00062 |
| Active (slope) | 0.011 | 0.0090 | [-0.00073 0.029] | 1.18 | 94 | 0.24 |

#### Random Effects Covariance Parameters

| Grouping Variable | Coefficient Names | Estimate | 95% CI |
|---|---|---|---|
| Participant (19 levels) | Control SD | 0.099 | [0.071 0.14] |
| Error | Residual SD | 0.044 | [0.038 0.052] |

#### Supplementary Analyses

##### Randomization procedures did not account for tVNS effects

We ruled out the possibility that our randomization procedure unintentionally favored tVNS effects through the ordering of stimulation conditions. Specifically, if a majority of participants completed tVNS after Sham conditions, any observed performance improvements during tVNS could be attributed to learning effects rather than tVNS-evoked effects. To examine this, we assessed the number of participants who underwent Sham before or after tVNS conditions in Experiments 2 and 3 (Table S7). If our randomization procedure favored tVNS effects, a majority of participants would have completed tVNS after Sham.

Excluding the conditioning period in Experiment 2, most participants completed taVNS 30 Hz and taVNS 3 Hz after Auricular Sham, indicating a potential for taVNS-related improvements. However, no significant taVNS effects were observed (Figure 4). Similarly, in Experiment 3, a majority of participants completed tcVNS 30 Hz after Arm stimulation, and taVNS 30 Hz after Auricular Sham. Yet, neither comparison revealed significant differences between conditions (Figure 2G and Figure 4F, respectively).

All significant tcVNS-evoked improvements were linked to either a balanced order of conditions (tcVNS 30 Hz vs Cervical Sham in Experiment 2; Figure 2C) or an unbalanced distribution that worked against tVNS effects (tcVNS 30 Hz vs tcVNS Sham in Experiment 3; Figure 2G). The only exception was that a majority of participants completed tcVNS triple pulse after Cervical Sham in Experiment 2. To examine whether this imbalance contributed to the significant visual acuity improvement (Figure 2F), we performed a resampling-based analysis to artificially balance the number of participants that completed Sham before and after tcVNS triple pulse (Figure S2). Five participants were sampled without replacement from the seven that completed tcVNS triple pulse after Cervical Sham. This resulted in a balanced dataset of 10 participants – 5 that completed tcVNS triple pulse before Cervical Sham and 5 that completed it after. Permutation paired t-tests compared sensory performance in this balanced dataset, with the resampling-based analysis repeated until each of the 21 possible combinations of 5 participants was assessed. P-values were corrected for multiple comparisons across all 21 iterations via the Max T method. Statistically significant tcVNS-evoked improvements were observed in each iteration (Figure S1). Therefore, the randomization procedure did not contribute to tcVNS-evoked sensory improvements in our study.

**Table S7.** Number of participants that completed tVNS conditions before or after Sham conditions. If the value in "tVNS after" exceeds the corresponding value in 'tVNS before', results could be biased toward observing tVNS effects. Bolded entries indicate comparisons that reached statistical significance.

| Experiment | Sham site | tVNS site | tVNS before | tVNS after |
|---|---|---|---|---|
| **2** | Auricular Sham | taVNS 30 Hz | 4 | 8 |
| | Auricular Sham | taVNS 3 Hz | 5 | 7 |
| | **Cervical Sham** | **tcVNS 30 Hz** | **6** | **6** |
| | **Cervical Sham** | **tcVNS triple-pulse** | **5** | **7** |
| **3** | Arm | taVNS 30 Hz | 6 | 6 |
| | Arm | tcVNS 30 Hz | 5 | 7 |
| | taVNS Sham | taVNS 30 Hz | 3 | 9 |
| | **tcVNS Sham** | **tcVNS 30 Hz** | **7** | **5** |

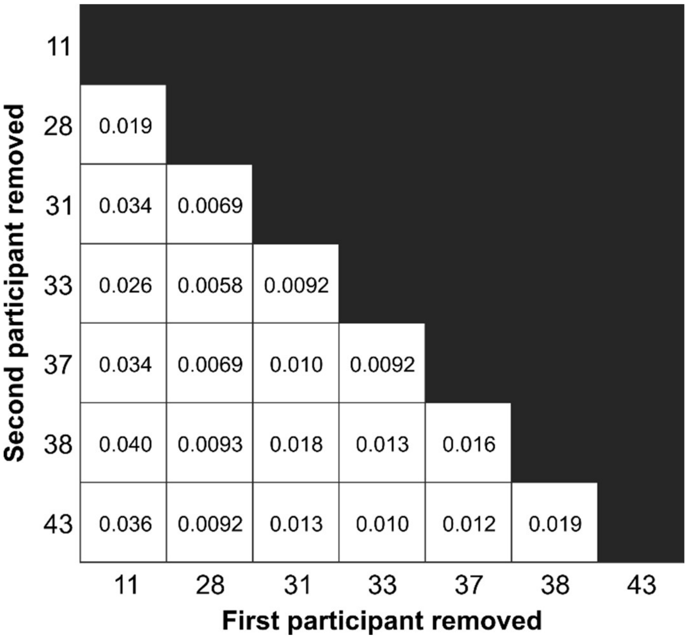


**Figure S1.** Statistically significant tcVNS-evoked visual acuity improvement unaffected by artificial balancing. Each cell in the heatmap depicts corrected p-values for permutation paired t-tests between visual performance during tcVNS triple pulse and Cervical Sham in Experiment 2. X- and y-axes labels indicates the participant ID removed during the artificial re-balancing procedure.
